# Stress-tolerant, recyclable, and autonomously renewable biocatalyst platform enabled by engineered bacterial spores

**DOI:** 10.1101/2022.03.16.484680

**Authors:** Yue Hui, Ziyu Cui, Seunghyun Sim

## Abstract

Here, we describe a stress-tolerant, recyclable, and autonomously renewable biocatalyst platform based on T7 RNA polymerase-enabled high-density protein display on bacterial spores (TIED). TIED uses high-level T7 RNA polymerase-driven expression of recombinant proteins specifically in sporulating cells to allow spontaneous assembly of recombinant fusion proteins on *B. subtilis* spore surface. TIED enables a high loading density in the range of 10^6^–10^7^ recombinant enzymes per spore, robust catalytic activities of displayed enzymes comparable to the respective free enzymes, and enhanced kinetic stability of displayed enzymes in methanol and elevated temperatures. Further, we demonstrate TIED-enzymes to be not only recyclable, but fully renewable after loss of activity through induction of germination and sporulation, enabling perpetual reuse of these immobilized biocatalysts.

**Graphical abstract:** 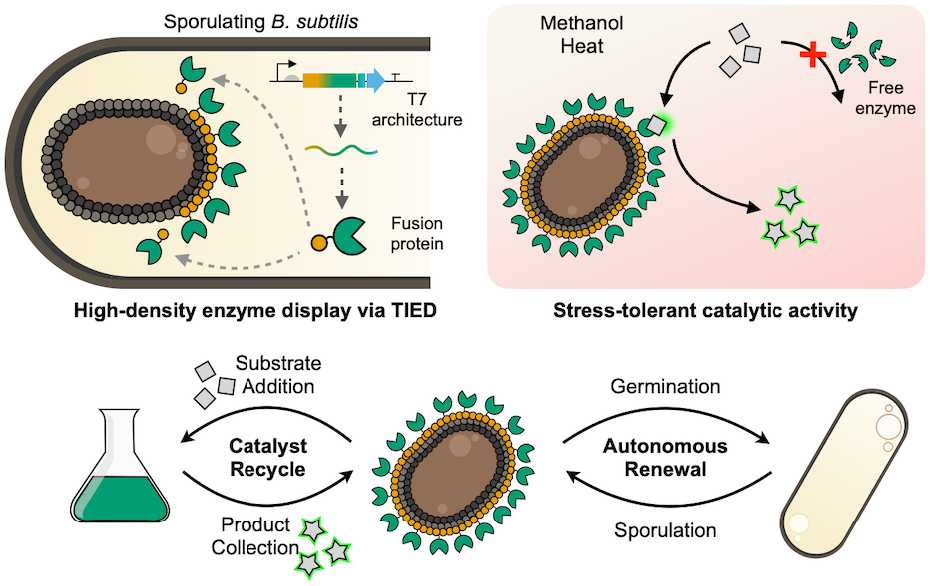

**Schematic illustration of the T7 RNA polymerase-enabled high-density protein display (TIED) on bacterial spores and its unique features as a biocatalyst platform**.

## INTRODUCTION

Enzymatic catalysis offers unique advantages for a wide range of chemical transformations because of its specificity, efficiency, and environmental friendliness.^1-4^ With advances in computational tools and directed evolution in recent years, enzyme engineering has expanded the portfolio of available biocatalysts beyond nature’s repertoire.^5-8^ Despite their apparent merits in chemical and pharmaceutical manufacturing processes, using enzymes in a scalable and economical manner remains challenging for a wide range of applications. In addition to costly purification, enzymes are less attractive in terms of stability and reusability compared to synthetic catalysts. Subjecting them to common reaction conditions, such as organic solvent or high temperature, often leads to denaturation and subsequent loss of function.^9,10^

Immobilizing enzymes on solid supports has been shown to alleviate these limitations and is considered essential for installing enzyme biocatalysts in large-scale production because it enables the recycling of enzymes.^11^ *In situ* immobilization of enzymes on bacterial spores features additional benefits by leveraging the autonomous biological process of protein synthesis and assembly on the spore surface. In particular, spores of *Bacillus subtilis* are well-suited for enzyme display. They can tolerate environmental stresses likely present in industrial settings, such as high temperature, nutrient deprivation, and organic solvents.^12^ In addition, a complex network of structural proteins comprising the outer layers of *B. subtilis* spores provides a plethora of possible anchoring motifs for enzyme loading.^13,14^ So far, a range of enzymes, including α-amylase, lipase, and β-galactosidase, have been installed on the surface of *B. subtilis* spores.^15-20^ However, despite a number of successful proof-of-concept studies, none so far have fully addressed major limitations of this system for practical applications: (1) low enzyme loading density, typically 10^2^–10^5^ enzymes per spore, leading to insufficient catalytic activity, and (2) loss of activity over time and in harsh conditions. Furthermore, quantitative and substrate-specific characterizations of catalytic behavior of spore-immobilized enzymes in comparison to their respective free-forms remain elusive.

This study demonstrates a novel biocatalyst platform that is stress-tolerant, recyclable, and autonomously renewable. This development was made possible by T7 RNA polymerase-enabled high-density protein display on bacterial spores (TIED) technology. We achieved 10^6^– 10^7^ enzymes per spore loading density with comparable catalytic performance to the respective free-form enzymes, enhanced catalytic activity in methanol, and increased temperature stability. In addition to being recyclable for multiple uses, enzyme-displaying spores can also be fully renewed upon attrition of their activity, allowing unlimited reuse of immobilized biocatalysts.

## RESULTS

### T7 RNA polymerase enabled high-density protein display on bacterial spores (TIED)

TIED combines the amplified expression of recombinant proteins by bacteriophage-derived T7 RNA polymerase (T7 RNAP), spore-specific promoters for activation of T7 RNAP, and spontaneous assembly of recombinant proteins on *B. subtilis* spore surface (Fig. 1a, left). We hypothesized that implementing T7 RNAP in the engineered genetic circuit designed to be active during late-stage sporulation would amplify the expression of fusion proteins, which subsequently assemble on the spore surface. We devised a panel of TIED constructs with three endogenous, σ_K_-specific promoters, *P*_*cotG*_, *P*_*cotV*_, and *P*_*cotZ*_. Target proteins were fused to the C-terminus of a coat protein, which assembles with other coat proteins to form the outer protein layers of *B. subtilis* spores. We chose the two morphogenic coat proteins suggested to play a crucial role in the assembly of the outermost crust layer, CotY and CotZ,^13,14,21,22^ for our design of TIED. An epitope tag, 3×FLAG, was incorporated on the C-terminus for immunolabeling and quantification of target proteins. To evaluate the performance of the TIED platform, we built genetic constructs with the endogenous promoters directly driving the expression of the fusion protein (Fig. 1a, right), analogous to the genetic architectures reported in previous studies.^15-20^ We selected a fluorescent reporter protein, mWasabi, as the first target to observe its overall expression level and spore localization in both TIED and endogenous expression systems.

**Fig. 1:**
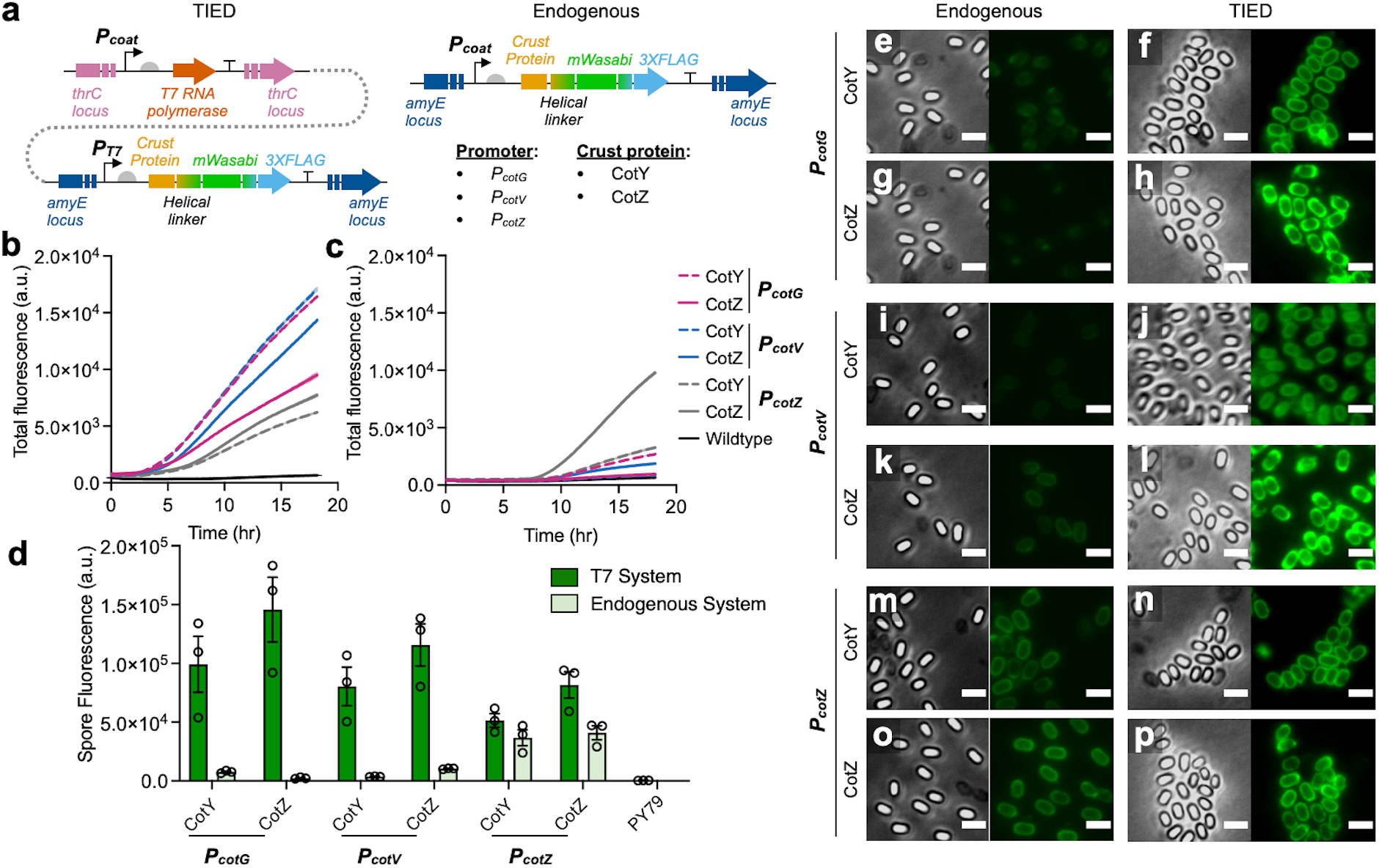
T7 RNA polymerase-enabled high-density protein display on *B. subtilis* spores (TIED). **a**, Genetic constructs of mWasabi display via TIED and endogenous constructs. Six constructs with combinations of three promoters (*P*_*cotG*_, *P*_*cotV*,_ and *P*_*cotZ*_) and two crust proteins (CotY and CotZ) were built for both systems. **b**, Total expression level of fluorescent fusion proteins in TIED-mWasabi variants. **c**, Total expression level of fluorescent fusion proteins in endogenous constructs. Each variant was grown to OD_600_ of 0.5 in LB and resuspended in an equal volume of SM medium. Fluorescence was continuously monitored after resuspension for 18 h. (*n* = 3 biological replicates). **d**, Spore fluorescence of the TIED variants, the endogenous counterparts, and their parental strain (PY79). Spores were harvested by lysozyme digestion (50 *μ*g mL^-1^) of the sporulating cells. Spores were washed twice and resuspended in an equal volume of PBS for fluorescence measurement. **e–p**, Representative phase-contrast and fluorescent microscopic images of spores of each variant. Fluorescence (green) of each sample was measured with 488 nm laser (100 ms exposure time, 3% laser, *n* = 3 biological replicates). See Extended Data Fig. 1c and 1d for images of the corresponding cells before lysozyme digestion. Scale bars, 2 *μ*m.

Upon inducing sporulation to *B. subtilis* cells, total expression levels of the fluorescent fusion protein were recorded for all 12 strains (Figs. 1b and 1c). In general, TIED constructs resulted in higher total expression of fluorescent fusion proteins and earlier activation compared to their endogenous counterparts. In particular, *P*_*cotG*_- and *P*_*cotV*_-driven TIED variants showed a more dynamic increase in fluorescence compared to *P*_*cotZ*_-driven ones (Fig. 1b). Meanwhile, *P*_*cotZ*_-driven constructs outperform the rest in the endogenous constructs (Fig. 1c). The difference in the promoter behavior of *P*_*cotZ*_ in TIED and endogenous counterparts may be attributed to the level of expression of T7 RNAP and fusion proteins. We speculate that the amount of T7 RNAP produced in *P*_*cotZ*_-driven TIED constructs is higher than others, creating a more burdensome and toxic environment for individual cells,^23,24^ which could lead to suboptimal output for fusion proteins.

We further investigated the distribution of the fluorescent fusion proteins through microscopic analysis and bulk fluorescence measurement of purified spore solutions. Despite the TIED system exhibiting a higher total expression level of the fluorescent fusion proteins (Fig. 1b), as forespores approach maturation, the fluorescence emission from the spore surface was significantly reduced, and most of the fluorescence emission was detected in the residual vegetative cells (Extended Data Fig. 1a). We hypothesized that a group of proteases active upon completion of mature spore formation to lyse the mother cell could have degraded fluorescent fusion proteins on the spore surface.^25^ Based on this hypothesis, we set out to improve the retainment of fusion proteins on spores by developing an early harvest protocol to collect mature spores while they are still inside their mother cells. Compared to natural maturation and spontaneous release from the mother cells, spores harvested with our protocol showed higher spore fluorescence in both endogenous and TIED constructs (Extended Data Figs. 1b, 1e, and 1f). Based on the bulk fluorescent intensities of the spore solutions normalized by the optical density at 600 nm (Fig. 1d) and microscopic analysis (Figs. 1e–1p), TIED generally outperformed their endogenous counterparts. In particular, TIED constructs with *P*_*cotG*_ and *P*_*cotV*_ promoters show expression levels two orders of magnitude higher than their endogenous counterparts as measured by fluorescence. In endogenous constructs, the expression level of the fluorescent fusion protein from the entire population has a positive correlation with the spore-specific fluorescence emission, suggesting that the total production of the fluorescent fusion protein is the determining factor for the number of fusion proteins on the spore surface (Extended Data Fig. 1h). In TIED, the expression level of the fluorescent fusion protein did not show any correlation with the spore-specific emission (Extended Data Fig. 1g), indicating the production yield of the fusion protein is not the determining factor for the spore display. We speculate that T7 RNAP-based expression of fusion proteins saturates the available translation machinery in sporulating cells. As a result, the interplay between the timing of the fusion protein expression, coat-protein assembly, and protease-mediated control of the late-stage sporulation determines the density of the displayed fusion proteins.

### Catalytic performance of TIED-LipB

Leveraging the TIED technology combined with an early harvest strategy, we devised a versatile platform for displaying recombinant proteins in high density on bacterial spores. After validating the performance of TIED with a fluorescent protein, we sought to display functional enzymes on *B. subtilis* spores (Fig. 2a). We chose *B. subtilis* lipase B (LipB) as our first target because of its relatively small size (19.5 kDa) and the general importance of lipases in industrial applications, including bulk production of food ingredients, detergents, and pharmaceuticals.^26^ Six TIED combinations with three promoters and the two anchor proteins resulted in phase-bright spores (Fig. 2b). Loading densities achieved by previous studies in spore-based enzyme display typically ranged from 10^2^ to 10^5^ enzymes per spore, based on enzymatic activities or dot blot experiments. Loading densities of LipB on each TIED construct were determined by quantitative western blot using spore lysate containing the crust and coat proteome. Among all TIED variants, *P*_*cotG*_ and CotZ fusion partner combination resulted in the highest loading density of LipB, 1.75 × 10^7^ LipB per spore (Fig. 2c, Extended Data Figs. 4a and 5a). Corresponding to the relative loading densities of the TIED constructs, normalized lipase activity assessed by colorogenic *p*-nitrophenol palmitate (C16) substrate conversion (Fig. 2d) shows that the *P*_*cotG*_ and CotZ combination exhibits the highest substrate conversion compared to all other TIED constructs by orders of magnitude (Fig. 2e). Subsequent enzymatic analyses of TIED-LipB, compared to the free-form LipB, were performed with this variant.

**Fig. 2:**
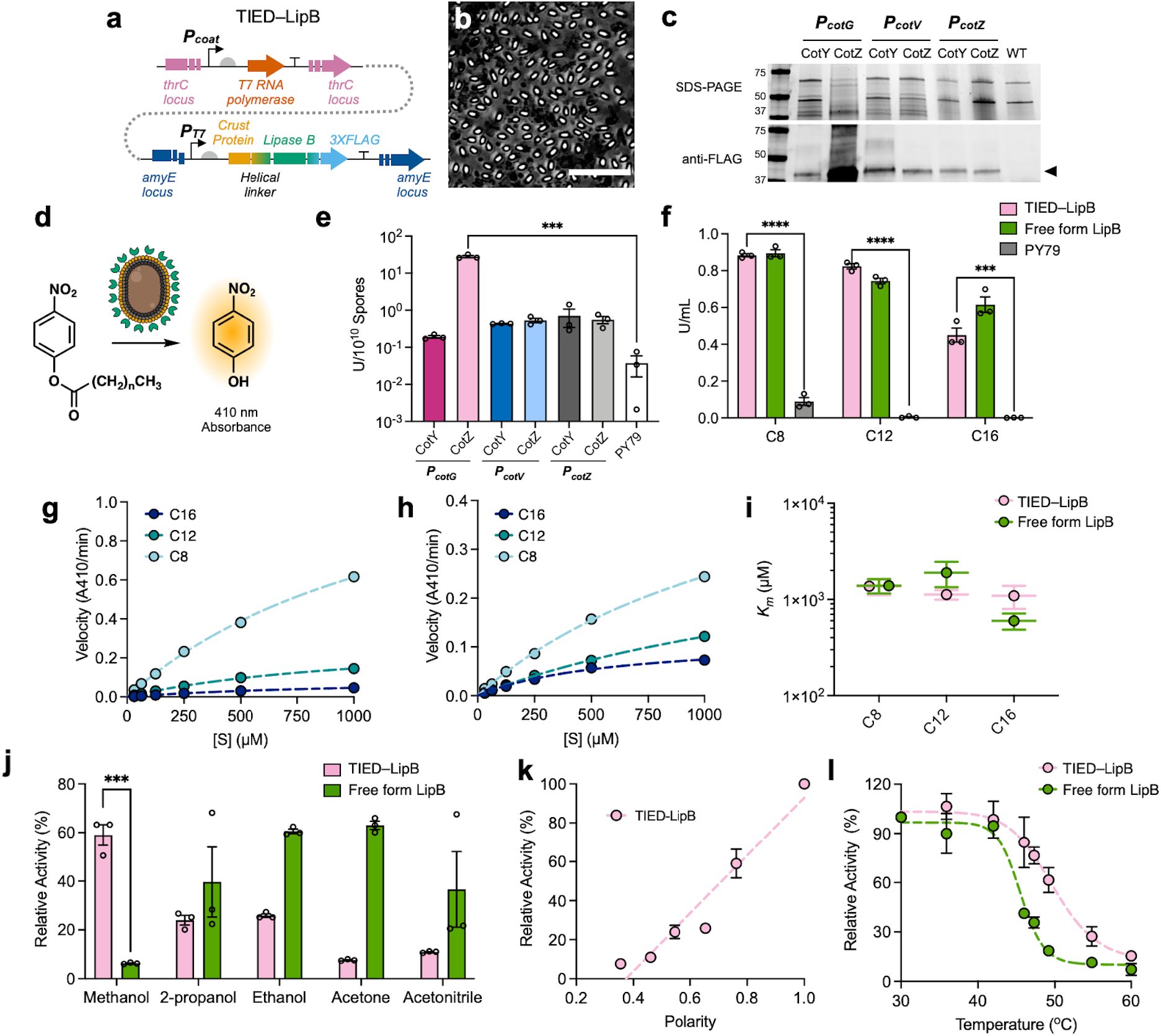
Catalytic performance of TIED-LipB. **a**, Genetic constructs of TIED-LipB. **b**, A representative phase-contrast image of TIED-LipB (with the combination of promoter *P*_*cotG*_ and CotZ fusion partner) after lysozyme digestion. Scale bar, 10 *μ*m. **c**, SDS-PAGE gel and western blot images of spore lysates of each variant and their parental strain PY79 (WT). Lysate of PY79 (WT) was included as a negative control. The protein content of each lysate, measured by BCA assay, was normalized to 20 *μ*g for loading in each lane. Resolved proteins were transferred for western blot analysis with primary anti-FLAG and secondary Alexa Fluor 647-conjugated antibodies. Bands corresponding to monomers of the respective fusion proteins were indicated by the black triangle. **d**, Schematic illustration of the colorimetric assay for TIED-lipase activity. **e**, Enzymatic activity of TIED-LipB variants and PY79 spores for hydrolysis of *p*-nitrophenol palmitate (C16, 1 mM). The enzyme activity unit, U, is defined as the conversion of 1 *μ*mol of a substrate into *p*-nitrophenol in 1 h at 42 °C. **f**, Normalized activity of TIED-LipB (combination of *P*_*CotG*_ and CotZ fusion partner) and free-form LipB for hydrolysis of *p*-nitrophenol palmitate (C16), *p*-nitrophenol dodecanoate (C12), and *p*-nitrophenol octanoate (C8). Spores of PY79 (OD_600_ of 0.5) analyzed with the same method were included as a negative control. **g**, Reaction velocities of TIED-LipB (combination of *P*_*CotG*_ and CotZ fusion partner, OD_600_ of 0.5) as a function of substrate concentrations. **h**, Reaction velocities of the free-form LipB (1 *μ*g mL^−1^) as a function of substrate concentrations. Dotted lines indicate fitting with the Michaelis Menten model (*n* = 3 biological replicates, see full kinetics data in Extended Data Fig. 6). **i**, The Michaelis Menten constant, *K*_*m*_, of TIED-LipB (combination of *P*_*CotG*_ and CotZ fusion partner) and the free-form LipB for hydrolysis of each substrate, determined from Michaelis Menten fitting. (Error bar indicates ± 90% confidence interval). **j**, Relative activities of TIED-LipB (combination of *P*_*CotG*_ and CotZ fusion partner, OD_600_ of 0.5) and the free-form LipB (100 *μ*g mL^−1^) for hydrolysis of C16 in organic solvents, normalized to their respective activity in Tris-HCl (pH 8.0) buffer. **k**, Relative activities of TIED-LipB (combination of *P*_*CotG*_ and CotZ fusion partner) as a function of solvent polarity. The dotted line indicates fitting with linear regression (*n* = 3 biological replicates). **l**, Relative activities of TIED-LipB (combination of *P*_*CotG*_ and CotZ fusion partner, OD_600_ of 0.5) and the free-form LipB (100 *μ*g mL^−1^) for hydrolysis of C16 at temperatures from 30 to 60 °C, normalized to their respective activities at 30 °C. The half inactivation temperature was determined by fitting a simple Boltzmann sigmoid function (*n* = 3 biological replicates). *P* values were determined by two-tail t-tests. \**P* < 0.05, \*\**P* < 10^−2^, \*\*\**P* < 10^−3^, \*\*\*\**P* < 10^−4^.

We investigated the effect of substrate bulkiness on the overall efficiency of TIED-LipB on spores with three substrates: C16, *p*-nitrophenol dodecanoate (C12), and *p*-nitrophenol octanoate (C8). Both TIED-LipB and the free-form LipB convert substrate with shorter alkyl chains more efficiently, and TIED-LipB showed hydrolysis activity for these substrates on-par to that of free-form LipB (Fig. 2f). We performed a traditional Michaelis-Menten kinetic study to evaluate the overall catalytic performance. Both TIED-LipB (Fig. 2g, Extended Data Fig. 6a) and the free-form LipB (Fig. 2h, Extended Data Figure 6b) showed a similar substrate-dependent, hyperbolic Michaelis-Menten kinetics with no apparent signs of substrate inhibition in the experimental concentration range. The apparent Michaelis-Menten constants (*K*_*M*_) of TIED-LipB and the free-form LipB were similar to each other for all substrates (Fig. 2i, Extended Data Figure 6c). Taken together, these results show the high efficiency and substrate binding affinity of the TIED-LipB to be comparable to free-form LipB.

We tested if the densely arranged, immobilized TIED-LipB exhibit better tolerance towards common stressors: organic solvents and elevated temperature. Polar organic solvents are substantially more destructive for enzyme activity because they penetrate and exchange tightly bound water from enzyme active sites.^27-29^ In particular, methanol deactivates lipases from various origins,^30-32^ which is a major limitation for industrially-relevant reactions, such as enzymatic biodiesel synthesis.^33^ We subjected TIED-LipB to a series of water-miscible and polar organic solvents.^34^ In contrast to the free-form LipB, to our surprise, a positive correlation between solvent polarity and relative enzymatic activity of TIED-LipB is observed (Figs. 2j and 2k, Extended Data Fig. 6d). Strikingly, TIED-LipB retains approximately 60% of catalytic activity in methanol, whereas the free-form LipB only showed 6% activity in the same condition (Figs. 2j). TIED-LipB and the free-form LipB likely experience different local environments – TIED-LipB is surface-bound and constitutes a densely packed protein layer, whereas the free-form LipB is in a bulk organic solvent. Solvent polarity, orientation, and solvated substrate anisotropy at the interface differ from those of the bulk solution and affect heterogeneous catalysis.^35-37^ We attribute this unique solvent polarity–activity trend to the heterogeneous nature and dense protein layers of the TIED system. In addition, TIED-LipB showed better temperature stability than the free-from LipB, increasing the half-inactivation temperature from 45 to 50 °C (Figs. 2l). Studies have shown that increasing the structural rigidity of enzymes is an effective strategy to improve the kinetic stability towards elevated temperatures.^33,38-40^ As such, the enhanced thermal stability is likely due to the increased rigidity of individual enzymes in densely packed protein layers on the spores. Taken together, these results demonstrate that TIED-LipB shows robust enzyme activity, comparable to the free-form LipB, and enables the use of LipB in traditionally harsh conditions.

### Catalytic performance of TIED-LipA

We selected lipase A (LipA), another extracellular lipase secreted by *B. subtilis* during vegetative growth, as our next target. Despite the high homology to LipB and similar α/β hydrolase core structure, LipA shows differences in substrate specificity, activity, and stability.^41^ Four TIED-LipA constructs were built, and each spore morphology was confirmed by phase-contrast optical microscopy (Figs. 3a, 3b, and Extended Data Fig. 2a). With the exception of the *P*_*cotG*_ and CotZ combination TIED construct, all other TIED combinations resulted in high levels of expression and assembly of LipA on the spore surface (Fig. 3c). The loading densities of the three successful TIED constructs are between 9.49 × 10^6^ and 2.93 × 10^7^ (Extended Data Figs. 4 and 5b-d). Consistent with their high loading density, the three TIED-LipA constructs all exhibited robust hydrolysis activity towards C16 substrate (Fig. 3d). We chose the TIED-LipA variant with the highest enzymatic activity (*P*_*cotV*_ and CotZ combination) for evaluating the enzymatic performance along with the free-form LipA. Enzymatic hydrolysis kinetics of TIED-LipA and the free-form LipA were measured and fitted to the Michaelis Menten model (Figs. 3e and 3f, Extended Data Figs. 7a and 7b). Similar to LipB, both TIED-LipA and the free-form LipA show faster kinetics for smaller substrates. Higher values of *K*_*M*_ in the case of TIED-LipA indicate that the substrate affinities to the enzyme are lower than those of the free-form LipA (Fig. 3g, Extended Data Fig. 7c). We attribute this trend to the local environment on the spore surface where the active site of immobilized LipA could be distorted due to orientational restrictions, resulting in less favorable substrate access. TIED-LipA also showed significantly higher activity in methanol compared to the free-form LipA (Fig. 3h) and a positive correlation between solvent polarity and enzyme activity (Fig. 3i). No such correlation was observed for the free-form LipA (Extended Data Figs. 7d). Furthermore, TIED-LipA exhibited superior thermal stability than the free-form LipA, with half inactivation temperature increased from 51 to 57 °C, likely due to the increased enzyme rigidity in the restricted local environment (Fig. 3j).

**Fig. 3:**
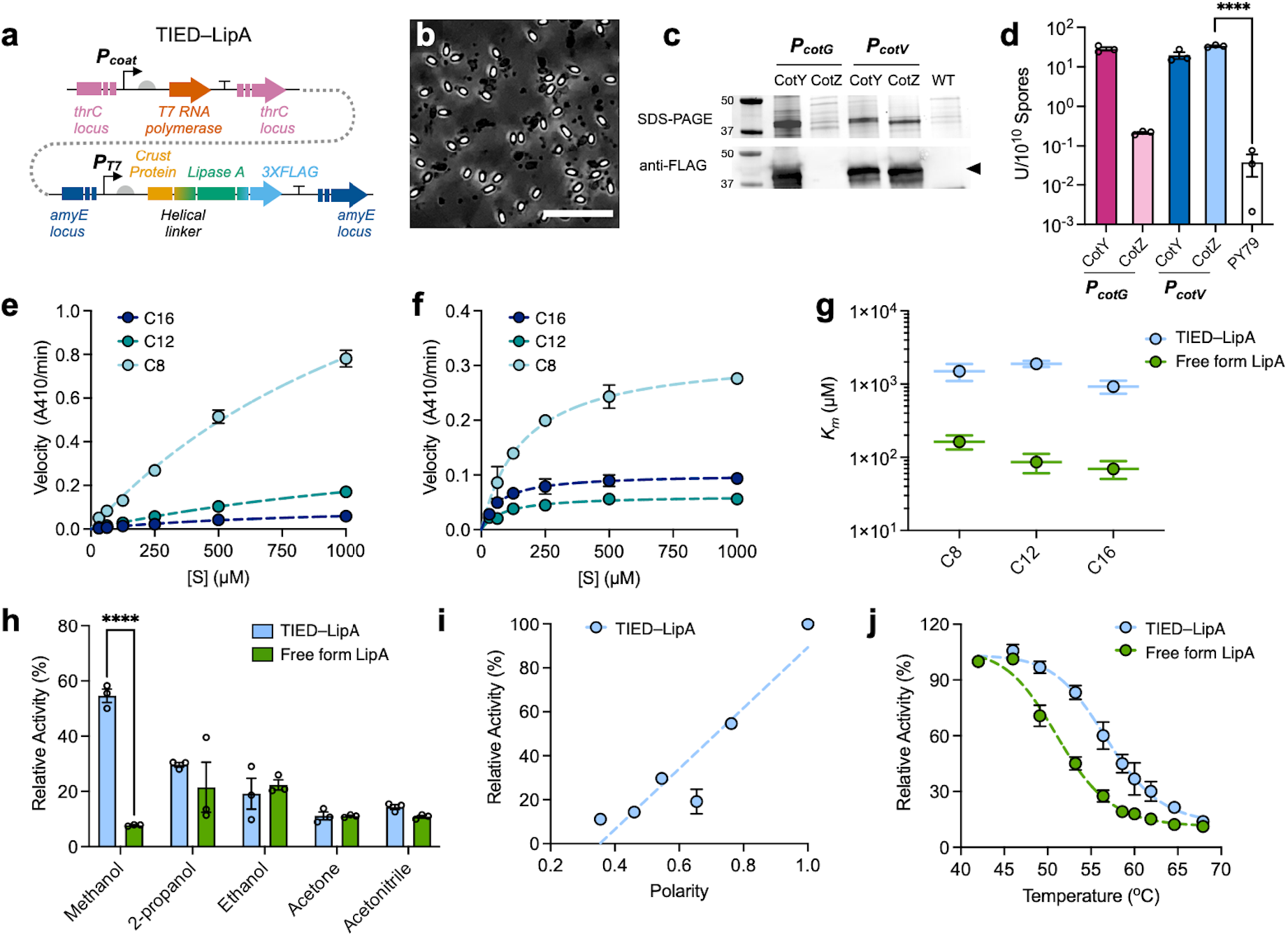
Catalytic performance of TIED-LipA. **a**, Genetic constructs of TIED-LipA. **b**, A representative phase-contrast image of TIED-LipA (with the combination of promoter *P*_*cotV*_ and CotZ fusion partner) after lysozyme digestion. Scale bar, 10 *μ*m. **c**, SDS-PAGE gel and western blot images of spore lysates of each variant and their parental strain PY79 (WT). Lysate of PY79 (WT) was included as a negative control. Protein content of each lysate, measured by BCA assay, was normalized to 20 *μ*g for loading in each lane. Resolved proteins were transferred for western blot analysis with primary anti-FLAG and secondary Alexa Fluor 647-conjugated antibodies. Bands corresponding to monomers of the respective fusion proteins were indicated by the black triangle. **d**, Enzymatic activity of TIED-LipA variants and PY79 spores for hydrolysis of *p*-nitrophenol palmitate (C16, 1 mM). The enzyme activity unit, U, is defined as the conversion of 1 *μ*mol of a substrate into *p*-nitrophenol in 1 h at 42 °C. **e**, Reaction velocities of TIED-LipA (combination of *P*_*CotV*_ and CotZ fusion partner, OD_600_ of 0.5) as a function of substrate concentrations. **f**, Reaction velocities of the free-form LipA (1 *μ*g mL^−1^) as a function of substrate concentrations. Dotted lines indicate fitting with Michaelis Menten model (*n* = 3 biological replicates, see full kinetics data in Extended Data Fig. 7). **g**, The Michaelis Menten constant, *K*_*m*_, of TIED-LipA (combination of *P*_*CotV*_ and CotZ fusion partner) and the free-form LipB for hydrolysis of each substrate, determined from Michaelis Menten fitting. (Error bar indicates ± 90% confidence interval). **h**, Relative activities of TIED-LipA (combination of *P*_*CotV*_ and CotZ fusion partner, OD_600_ of 0.5) and the free-form LipA (100 *μ*g mL^-1^) for hydrolysis of C16 in organic solvents, normalized to their respective activity in Tris-HCl (pH 8.0) buffer. **i**, Relative activities of TIED-LipA (combination of *P*_*CotV*_ and CotZ fusion partner) as a function of solvent polarity. The dotted line indicates fitting with linear regression (*n* = 3 biological replicates). **j**, Relative activities of TIED-LipA (combination of *P*_*CotV*_ and CotZ fusion partner, OD_600_ of 0.5) and the free-form LipA (100 *μ*g mL^−1^) for hydrolysis of C16 at temperatures from 42 to 68 °C, normalized to their respective activities at 42 °C. The half inactivation temperature was determined by fitting a simple Boltzmann sigmoid function (*n* = 3 biological replicates). *P* values were determined by two-tail t-tests. \**P* < 0.05, \*\**P* < 10^−2^, \*\*\**P* < 10^−3^, \*\*\*\**P* < 10^−4^.

### Catalytic performance of TIED-APEX2

To demonstrate the capability of TIED technology to synthesize and immobilize heterologous enzymes that are non-native to *B. subtilis*, we chose APEX2, an engineered ascorbate peroxidase, as our last target. In addition to extensive applications in proximity-based protein labeling and electron microscopy,^42^ APEX2 has recently been shown to enable conductive polymer synthesis *in vivo*.^43^ Analogous to TIED-LipB and TIED-LipA, all TIED-APEX2 constructs (Fig. 4a) yield typical spore formation (Fig. 4b, and Extended Data Fig. 2). Western blot analysis of the spore coat and crust proteome revealed the presence of full-length APEX2 in all TIED-APEX2 variants, except for the *P*_*cotZ*_ and CotZ combination (Fig. 4c). The lower molecular weight product observed in this combination indicates possible protease-mediated cleavage of the fusion protein. The loading densities of the full-length APEX2 in the successful TIED-APEX2 constructs, measured by quantitative western blot analyses, range from 2.85 × 10^6^ to 1.71 × 10^7^ (Extended Data Figs. 4 and 5e-h). In the presence of H_2_O_2_, APEX2 rapidly converts Amplex Red substrate into a fluorogenic product, resorufin (Fig. 4d). Corresponding to the western blot results, TIED-APEX2 spores with a high loading density (> 10^6^ enzymes per spore) all showed a robust enzymatic activity as probed by the Amplex Red conversion (Figs. 4e and 4f). In addition, kinetic analysis of these TIED-APEX2 variants revealed *K*_*M*_ similar to the free-form APEX2, indicating a similar substrate affinity (Figs. 5g and 5h, and Extended Data Fig. 8). These results demonstrate that TIED technology can effectively accommodate target enzymes that are completely foreign to *B. subtilis*.

**Fig. 4:**
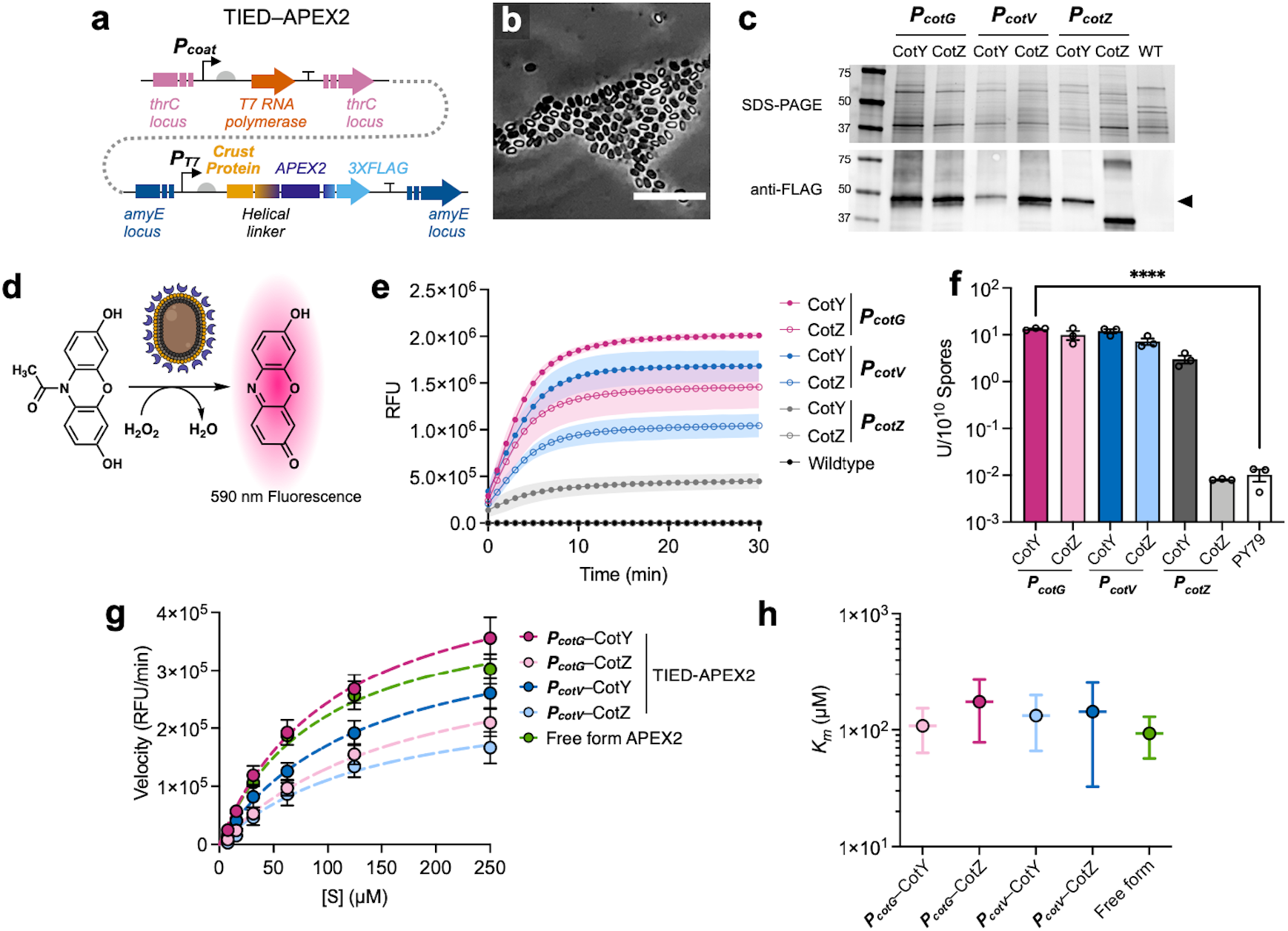
Catalytic performance of TIED-APEX2. **a**, Genetic constructs of TIED-APEX2. **b**, A representative phase-contrast image of TIED-APEX2 (with the combination of promoter *P*_*cotG*_ and CotY fusion partner) after lysozyme digestion. Scale bar, 10 *μ*m. **c**, SDS-PAGE gel and western blot images of spore lysates of each variant and their parental strain PY79 (WT). Lysate of PY79 (WT) was included as a negative control. The protein content of each lysate, measured by BCA assay, was normalized to 20 *μ*g for loading in each lane. Resolved proteins were transferred for western blot analysis with primary anti-FLAG and secondary Alexa Fluor 647-conjugated antibodies. Bands corresponding to monomers of the respective fusion proteins were indicated by the black triangle. **d**, Schematic illustration of the TIED-APEX2 catalyzed conversion of Amplex Red to resorufin. **e**, Fluorescence time trace resulting from resorufin formation catalyzed by TIED-APEX2 variants and PY79. **f**, Enzymatic activity of TIED-APEX2 variants and PY79 spores for conversion of Amplex Red (125 *μ*M). The enzyme activity unit, U is defined as the conversion of 1 *μ*mol of Amplex Red to resorufin in 1 h at 25 °C. **g**, Reaction velocities of TIED-APEX2 variants (OD_600_ of 0.5) and the free-form APEX2 (1 *μ*g mL^−1^) as a function of substrate concentrations. Dotted lines indicate fitting with Michaelis Menten model (*n* = 3 biological replicates, see full kinetics data in Extended Data Fig. 8). **h**, The Michaelis Menten constant, *K*_*m*_, of TIED-APEX2 variants and the free-form APEX2 for Amplex Red conversion, determined from Michaelis Menten fitting. (Error bar indicates ± 90% confidence interval). *P* values were determined by two-tail t-tests. \**P* < 0.05, \*\**P* < 10^−2^, \*\*\**P* < 10^−3^, \*\*\*\**P* < 10^−4^.

### Recycling and Autonomous Renewal of TIED-Enzymes

The design of the TIED system allows recycling and autonomous renewal of enzymes to fully recover original activity on-demand, enabling theoretically infinite reuse of biocatalysts (Fig. 5a). We first set out to test the recyclability of TIED-enzymes, which is one of the major known advantages of enzyme immobilization. After each reaction cycle, TIED-LipA was immediately recollected by centrifugation and subjected to the next cycle. With C8 and C12 substrates, TIED-LipA maintains almost 100% activity for at least 20 cycles (Extended Data Figs. 9a and 9b). In the case of C16 substrate, the enzymatic conversion diminished to below 50% after 10 rounds of recycling. Similar to TIED-LipA, conversion of C16 by TIED-LipB declined after 5 rounds of recycling (Extended Data Fig. 9c). TIED-APEX2 retained 83.6% of its original activity after 10 consecutive cycles (Extended Data Fig. 9d).

Another obvious merit of the TIED system is that the synthesis and assembly of recombinant enzymes are completely autonomous. Therefore, once the activity is reduced to a critical point, TIED enzymes can be fully renewed by germinating spores into cells, growing cells, and subsequently triggering sporulation. We demonstrated this novel concept with TIED-LipA for conversion of C16, the substrate that causes attrition of enzymatic activity after 10 cycles. After completion of the first 10 cycles, TIED-LipA spores (Figs. 5b) were germinated in nutrient-rich media (Figs. 5c–e), after which sporulation was induced by media shift. The catalytic performance of renewed TIED-LipA in cycle 11 is fully comparable to the pristine TIED-LipA in cycle 1 (Fig. 5g). Renewals after consecutive 10 cycles of TIED-LipA (cycle 21 and cycle 31) all result in full recovery of activity (Fig. 5g). Similar to TIED-LipA, TIED-LipB also recovered its full activity upon renewal after cycle 6, allowing multiple recycling and renewal rounds (Extended Data Fig. 9c). Recycling and renewal ability of TIED-LipA was observed even in methanol, albeit with a lower enzymatic conversion and more rapid loss of activity than in aqueous solutions (Fig. 5h). Multiple renewals of TIED-LipA after 5 consecutive cycles result in full recovery of enzymatic activity, which is consistent with the known tolerance of *B. subtilis* spores against organic solvents.

**Fig. 5:**
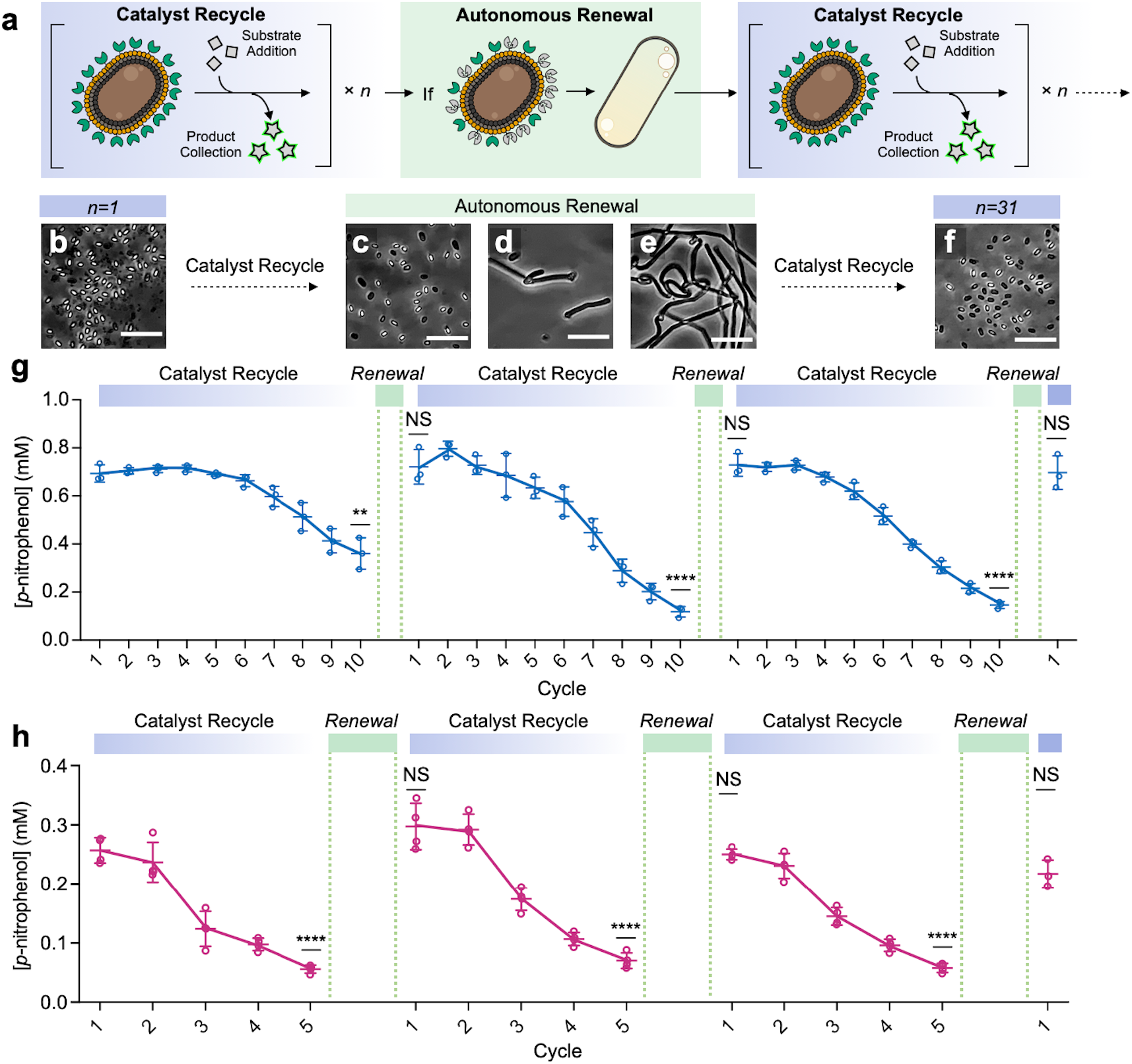
Recycling and Autonomous Renewal of TIED-Enzymes. **a**, Schematic illustration of recycling and autonomous renewal of TIED-enzymes. **b**, A representative phase-contrast microscopic image of TIED-LipA (combination of *P*_*cotV*_ and CotZ fusion partner) spore after the 1^st^ cycle of reaction with *p*-nitrophenol palmitate (C16, 0.8 mM) in Tris-HCl buffer (pH 8.0). **c**, A representative phase-contrast microscopic image of TIED-LipA (combination of *P*_*cotV*_ and CotZ fusion partner) spore germination in LB medium after 2 h. **d**, A representative phase-contrast microscopic image of TIED-LipA (combination of *P*_*cotV*_ and CotZ fusion partner) spore germination in LB medium after 3 h. **e**, A representative phase-contrast microscopic image of TIED-LipA (combination of *P*_*cotV*_ and CotZ fusion partner) spore germination in LB medium after 4 h. **f**, A representative phase-contrast microscopic image of TIED-LipA (combination of *P*_*cotV*_ and CotZ fusion partner) spore after the completion of the 31^st^ cycle of reaction with C16 (0.8 mM) in Tris-HCl buffer. Scale bar, 10 *μ*m **g**, Recycling and renewal of TIED-LipA (combination of *P*_*cotV*_ and CotZ fusion partner, OD_600_ = 1) spores catalyzing the conversion of C16 (0.8 mM) in Tris-HCl buffer. Each reaction cycle lasts 10 min (*n* = 3 biological replicates). **h**, Recycling and renewal of TIED-LipA (combination of *P*_*cotV*_ and CotZ fusion partner, OD_600_ = 1) spores catalyzing the conversion of C12 (0.8 mM) in methanol. Each reaction cycle lasts 20 min (*n* = 4 biological replicates). *P* values were determined by two-tail t-tests against the pristine activity of TIED-LipA (1^st^ cycle). \**P* < 0.05, \*\**P* < 10^−2^, \*\*\**P* < 10^−3^, \*\*\*\**P* < 10^−4^.

## DISCUSSION

We developed TIED technology to immobilize enzymes in high density on bacterial spores leveraging a T7 RNAP expression system active specifically during sporulation. The versatility and utility of this platform are demonstrated with three different enzymes – two endogenous lipases from *B. subtilis* and an engineered peroxidase APEX2 – all exhibiting efficient substrate conversion comparable to the respective free enzymes. Compared to the previously reported spore display systems, TIED significantly enhances the loading capacity of recombinant enzymes (10^6^–10^7^ per spores). Enhanced thermal stability of displayed lipases was observed. In addition, TIED-lipases show high activity in methanol and a unique linear relationship between enzymatic activity and solvent polarity. This feature makes TIED technology desirable for practical applications, such as biodiesel synthesis. Departing from all previously reported enzyme immobilization strategies, TIED-enzymes are not only recyclable, but are fully renewable. The high efficiency from the high loading density, improved stability in harsh conditions, recyclability, and renewability of the TIED system compare favorably to existing enzyme immobilization technologies, making TIED a powerful and sustainable biocatalyst platform.

Our current work lays the foundation for expanding the scope of the TIED technology. Because of the highly interconnected nature of spore coat proteins,^15^ the maximum level of one anchor protein may be limited by the availability of their interaction partners. For example, co-expression of fusion proteins harnessing a well-known interaction pair, CotY and CotZ,^44,45^ may further enhance the loading capacity. In addition, different local environments that individual coat proteins reside in may have unexpected implications on enzyme thermostability and activity in organic solvents. With more varieties and combinations of anchor proteins, TIED can potentially offer a higher loading density and meet requirements for specific applications for end-users. Moreover, future generations of TIED will enable multifunctional biocatalysts by displaying more than one type of enzyme. Engineering enzymatic cascades on a single spore surface could be pursued by displaying all involved enzymes in close proximity. Another useful application is to display enzymes having different substrate preferences on a single spore. This new version of TIED will enable a fast and convenient approach for generating one-for-all biocatalysts useful for multiple chemical transformations.

While in this work we compared the performance of TIED to free-form enzymes, it is worth noting additional advantages of spore-surface display over cell-surface display. Genetically encoding heterologous enzymes usually reduces growth rate, viability, and cell density relative to parental nonproducing cells, which gives a strong selection pressure for mutations eliminating the burden. In the TIED system, burdensome expression of genetically-encoded fusion proteins occurs only after *B. subtilis* cells have committed to sporulation. Because the burden and cellular growth are decoupled in the TIED system, the selective pressure for mutations debilitating the enzymatic functionality is largely or entirely mitigated. This evolutionary stability is another strong merit for the TIED system as it allows for the unlimited renewal of biocatalysts. Overall, we believe that TIED technology will generate a large portfolio of biocatalysts useful in various practical applications and advance the field of sustainable catalysis.

## METHODS

### Design and cloning of genetic constructs in *B. subtilis*

Gene sequences of lipase A, lipase B, CotY, and CotZ were obtained from gene *estA, estB, cotY*, and *cotZ*, respectively, from the genome of *B. subtilis* PY79. The secretion signal sequences of estA and estB were excluded. Gene sequences of mWasabi and APEX2 were codon-optimized (IDT Codon Optimization Tool) based on their protein sequences.^42,46^ Gene sequences of T7 RNAP and the T7 promoter, along with the lac operator and hammered Ribozyme were amplified from a previously reported construct (GenBank MN005204).^47^ The endogenous promoters, *P*_*cotG*_, *P*_*cotZ*_, and *P*_*cotV*_, were amplified from the genome of *B. subtilis* PY79. A strong synthetic ribosomal binding site (RBS), MF001, was inserted between the promoter region and start codon.^47^ Between the target protein and the C terminus of CotY/CotZ, an 11-amino-acid spacer that forms a stable α-helical structure was placed.^48^ Three repeats of the FLAG tag were introduced to the C terminus of the fusion construct for immunostaining.

PCR-amplified parts of DNA were assembled using Golden Gate Assembly and inserted into plasmid backbones (New England Biolabs).^49^ The T7 driven gene circuits for fusion protein expression were inserted into vector PBS1C,^50^ which carries homology sites of *amyE* locus and a chloramphenicol resistance selection marker. The T7 RNAP expression circuit was cloned into PBS4S backbone,^51^ which contains the spectinomycin resistance gene, flanked by sequences homologous to the *thrC* locus. The resultant plasmids were transformed into *E. coli* DH10B (New England Biolabs, C3019) through electroporation and selected by carbenicillin resistance (100 *μ*g mL^-1^). Clones carrying plasmids with correct sequences were isolated for DNA extraction (QIAGEN, 56604785).

Purified plasmids were linearized by restriction enzyme for the transformation of *B. subtilis*. Chromosomal integration into *B. subtilis* PY79 genome via double cross-over was performed according to the reported two-step method.^52^ The resulting *B. subtilis* clones were selected on lysogeny broth (LB) agar plates supplemented with chloramphenicol (5 *μ*g mL^-1^) or (and) spectinomycin (100 *μ*g mL^-1^). To generate strains carrying TIED constructs with engineered circuits in two loci, *B. subtilis* PY79 was transformed first with the T7-driven fusion protein expression circuit into the *amyE* locus. Competence was induced in selected clones for the subsequence transformation of the construct for T7 RNAP expression.

The construct for expressing free enzymes contained a fusion protein sequence with a 6×His tag appended to the N terminus of the genes encoding lipase A, lipase B, or APEX2 and three repeats of FLAG tag on the C terminus. The assembled DNA-of-interest was inserted into a pET22b expression vector containing a T7 promoter and lac operon (Novagen). Ligated plasmids were electroporated into *E. coli* BL21 (DE3) competent cells (Invitrogen). Clones were selected on LB agar plates supplemented with carbenicillin (100 *μ*g mL^-1^). The complete list of plasmids used and created in this study is provided in Supplementary Table 1.

### Bacterial growth and sporulation

Sporulation of wild-type and mutant *B. subtilis* strains was induced by the resuspension method.^53^ Starter cultures of *B. subtilis* cells were grown till saturation in LB medium. The starter culture was diluted 1:100 into a 50 mL fresh LB medium in a 250 mL Erlenmeyer flask and allowed to grow at 37 °C until the optical density at 600 nm (OD_600_) reached 0.5. Culturing time for different variants depended on their respective growth rate and was optimized for each. Cells were harvested by centrifugation (3000 g, 8 min), and the cell pellets were resuspended in an equal volume of SM medium. The resuspended cells were transferred back to Erlenmeyer flasks and grown for 14 h with agitation (37 °C, 250 rpm). The resulting cells were collected (4000 g, 10 min) and treated with 50 *μ*g mL^-1^ lysozyme (Sigma-Aldrich, L6876) in phosphate buffered saline (PBS, pH 7.2, gentrox, 30-025). The harvested spores were washed with PBS twice and checked for morphology using ECHO Revolve Microscope (inverted).

### Fluorescent microscopy and image analysis

2 *μ*L of TIED-mWasabi spore suspension in PBS were dropped on a microscopic slide, topped with a cover glass, and mounted on an ECHO Revolve Microscope (inverted mode). A 60x phase contrast lens and a 488 nm laser line (100 ms exposure time, 3% laser) were used for the imaging of spores. Images were exported and processed with Image J (NIH).

### Protein expression and purification

For the expression of the free-form LipA, LipB, and APEX2, the respective expression strain of *E. coli* was inoculated in 10 mL of LB, supplemented with 100 *μ*g mL^-1^ carbenicillin, and grew for 14 h at 37 °C. The saturated starter culture was then diluted into 500 mL Terrific Broth (Sigma-Aldrich) in 2800 mL baffled Erlenmeyer flask and allowed to grow at 37 °C (200 rpm) till OD_600_ of reached 0.5. The culture was cooled on ice for 10 min and induced with 1 mM IPTG. The induced culture was grown at 25 °C for 20 h (120 rpm). Cells were pelleted and resuspended in lysis buffer (50 mM NaH_2_PO_4_, 300 mM NaCl, 10 mM imidazole, pH 8.0) supplemented with 1 *μ*g mL^-1^ lysozyme, and incubated on ice for 1 h. The cellular structure was disrupted by sonication (Qsonica, Q125, 5s on, 5s off, 5 min, 100% amplitude), and the debris was pelleted by centrifugation (40,000 g, 1 h, 4 °C) and discarded. The lysate (20 mL) was incubated with 10 mL of Ni-NTA agarose (QIAGEN, 30230) for 30 min at room temperature. The protein-bound agarose resin was washed with 60 mL of wash buffer (50 mM NaH_2_PO_4_, 300 mM NaCl, 20 mM imidazole, pH 8.0) and eluted with 20 mL of elution buffer (50 mM NaH_2_PO_4_, 300 mM NaCl, 250 mM imidazole, pH 8.0). The eluted fraction was analyzed by SDS-PAGE to confirm the protein size and purity. Purified proteins were then flash-frozen at -80 °C and lyophilized for 48 h.

### Lipase activity assay

Spores with TIED-lipases were harvested by lysozyme treatment and resuspended in Tris-HCl buffer (50 mM pH 8.0, supplemented with 0.3% Triton-X). The lipase activity assay was conducted based on previously established protocols.^54^ 10 mM stock solution of lipase substrates, C8, C12, and C16 were prepared in isopropanol. Enzymatic activities ofTIED variants were evaluated by reacting spore suspension normalized to OD_600_ of 0.5 with 1 mM substrate C16 in 1 mL total volume for 1 h at 42 °C with agitation (800 rpm). After the reaction, spores were spun down, and 100 *μ*L of the supernatant containing the reaction product was loaded onto a 96-well microplate. The absorbance of each well at 410 nm was measured by a microplate reader (BioTek Synergy H1 Hybrid Multi-Mode, gain = 100). To convert the absorbance at 410 nm to product concentration, a calibration curve of *p*-nitrophenol concentration versus absorbance was plotted using solutions prepared with a standard compound (Sigma-Aldrich) (Extended Data Fig. 3b). The enzyme activity unit, U, is defined as the conversion of 1 *μ*mol of a substrate into *p*-nitrophenol in 1 h at 42 °C.

To compare the activity of TIED-LipB to free form LipB, spore displayed enzyme concentration (using 1 mL spore suspension of OD_600_ = 0.3) was calculated from quantitative western blot analysis. Purified and lyophilized free form LipB was dissolved in Tris-HCl buffer (50 mM pH 8.0, supplemented with 0.3% Triton-X) to match the enzyme concentrations of the spore suspensions. With each of the three substrates, reactions were performed at 42 °C for 1 h.

The kinetics assay was performed on a microplate reader. Each of the three substrates was serially diluted to concentrations ranging from 10 mM to 71.825 *μ*M in isopropanol. Spore suspension (OD_600_ = 0.5) was transferred to a 96-well microplate and pre-incubated at 42 °C for 10 min. Substrates of varying concentrations were then simultaneously diluted 1:10 to a final volume of 100 *μ*L. The absorbance at 410 nm in each well was continuously measured till the absorbance readouts plateaued.

### Lipase activity assays in organic solvents

For TIED-lipases, spores in 1 mL PBS (OD_600_ = 0.5) were pelleted and placed in a vacuum desiccator for 3 h to remove residual water. Dried spores were suspended in the respective organic solvents by brief sonication (VWR Symphony Ultrasonic Cleaner). Stock solutions of lipase substrates C8, C12, and C16 were diluted 1:10 into spore suspensions to a final concentration of 1 mM in 1 mL total volume. Lyophilized free-form enzymes (100 *μ*g mL^-1^) were directly dissolved in the respective solvents containing 1 mM substrates. Reactions were performed at 42 °C for 1 h, followed by centrifugation (10^4^ g, 10 min). The supernatants containing the reaction product were collected and heated at 80 °C for 10 min to deactivate any carried-over enzymes. Solvents in the supernatant were completely evaporated by a rotary evaporator (Buchi R-100, < 30 kPa), and the dried residues were dissolved in an equal volume of Tris-HCl buffer for colorimetric quantification.

### Thermal stability of Lipases

Temperatures used in the assays were implemented by a user-defined gradient protocol on a thermocycler (Bio-Rad, C1000). TIED-lipases (spore OD_600_ = 0.5) and the free-form lipases (100 *μ*g mL^-1^) dissolved in Tris-HCl buffer were preincubated at each temperature for 5 min to reach equilibrium. Reactions (total volume of 100 *μ*L) were initiated by spiking in C16 substrate to the final concentration of 1 mM and allowed to proceed for 1 h. Supernatants were collected for colorimetric measurements. Activities of LipA and LipB at elevated temperatures were determined relative to those at their respective optimal temperatures, 42 °C for LipA and 30 °C for LipB.

### APEX2 activity assay

Amplex Red reagent (Invitrogen, A12222) was dissolved in DMSO (≥ 99.7%) to prepare a 10 mM stock solution. Reactions were performed in a 96-well plate with 100 *μ*L total volume. Spores with TIED-APEX2 were suspended in PBS (pH 7.2). H_2_O_2_ was added from a 20 mM stock solution to the final concentration of 1 mM. The mixtures with spore (OD_600_ = 0.1) were transferred onto a microplate. The enzymatic reaction was initiated by adding Amplex Red stock solution to a final concentration of 125 *μ*M in each well, followed by manual mixing and immediate fluorescence monitoring (λ_ex_ = 530 nm, λ_em_ = 590 nm) for 1 h. The enzyme activity unit, U, is defined as the conversion of 1 *μ*mol of Amplex Red to resorufin in 1 h at 25 °C. For the kinetics assay, the Amplex Red stock solution was serially diluted to concentrations ranging from 10 mM to 71.825 *μ*M in DMSO. Substrates with varying concentrations were added 1:10 to each well containing TIED-APEX2 (in PBS with 1 mM H_2_O_2_). Fluorescence was continuously monitored under constant shaking till the emission readouts plateaued. To convert the fluorescence emission (RFU) to product concentration, a calibration curve of resorufin concentrations versus RFU was plotted using solutions prepared with a standard compound (Sigma-Aldrich, 635-78-9) (Extended Data Fig. 3c).

### Preparation of spore lysate and western blot analysis

For quantification of TIED-enzyme concentrations, concentrated spore suspension in PBS was serially diluted, and OD_600_ of each dilution was measured. Spores were collected and resuspended in an equal volume of a decoating buffer (100 mM NaOH, 100 mM NaCl, 100 mM dithiothreitol, and 10 g L^-1^ sodium dodecyl sulfate).^55^ Spores were decoated for 30 min at 65 °C. After centrifugation (10000g, 15 min), clear lysate containing solubilized spore coat and crust proteome was collected for western blot analysis. For comparing expression levels in different TIED variants, the protein concentration of each lysate sample was measured by BCA Protein Assay Kit (Thermo Scientific, 23250) according to the manufacturer’s manual. The original lysate was diluted 1:20 in PBS to lower the concentration of sodium dodecyl sulfate to below 5 mM. Samples with normalized protein content were loaded into each lane (20 *μ*g/lane) and resolved by SDS-PAGE (Bio-Rad, 456809). Total protein contents were imaged with the Bio-Rad Chemidoc MP Imaging System, and the gels were transferred to a PVDF membrane (Bio-Rad, 1620174) using a western blot transfer system (Trans-Blot Turbo Transfer System, 1.5 mm Gel Protocol). Membranes were blocked with 3% (w/v) BSA (Sigma-Aldrich, A9418) in 0.1% Tween 20 in phosphate buffered saline (PBST) at room temperature for 1 h to prevent non-specific labeling. Primary anti-FLAG antibody (1: 5000, Sigma Aldrich, F1804) was added, and the labeling was allowed to proceed for 1 h at room temperature. Membranes were then washed three times in PBST to remove unbound antibody and incubated with Alexa Fluor 647 conjugated secondary antibody (1: 2000, Goat anti-Mouse IgG (H+L) Highly Cross-Adsorbed Secondary Antibody, Alexa Fluor 647, Invitrogen, A21236) in PBST for 30 min in dark. Stained membranes were rinsed three times in PBST and imaged with Bio-Rad Chemidoc MP Imaging System. Fluorescent signals from each membrane were collected using the Cy5 channel (5 s exposure time, gain = 1), and the resulting images were subjected to the same image processing method with Bio-Rad Image Lab Touch Software to ensure a fair comparison.

### Quantification of TIED-enzyme loading density

Western blot images were processed with Image J (Extended Data Fig. 4).^56^ Different spore concentrations were used to prepare lysates in each lane. Intensities of all bands within the lane were plotted using an existing function in Image J (Plot Lanes). The area-under-peaks corresponding to the TIED-enzymes were segmented and quantified. To quantify the enzyme loading density in each TIED variant, the intensities of the bands corresponding to the monomer of the fusion protein were plotted against the OD_600_ of spore suspension used to generate the lysate. The linear range was manually determined and fitted with linear regression (Extended Data Fig. 5). Colony-forming units (CFU) corresponding to each OD_600_ were determined from a CFU-spore OD_600_ calibration curve. Purified spores were diluted in PBS to OD_600_ between 0.1 to 1. Each sample was serially diluted and plated on LB agar plates, and the CFUs were calculated based on the number of colonies on the plates. Linear regression was then fitted to the data (Extended Data Fig. 3a). For each target protein, free enzyme solutions ranging from 1 mg mL^-1^ to 1.56 *μ*g mL^-1^ were prepared with purified and lyophilized protein powder. After SDS-PAGE and subsequent western blot analysis, intensities of the bands corresponding to a specific concentration of free enzymes were quantified. A calibration curve of band intensities versus enzyme concentrations was plotted for each target enzyme. The linear range was determined in each case and fitted with regression (Extended Data Fig. 5). Together with the OD_600_ and CFU conversion plot, TIED-enzyme loading densities for each variant were determined with these calibration curves.

### Recycle and renewal of spore with TIED-enzymes

For aqueous phase reactions, spores with TIED-LipA were suspended in 50 mM Tris-HCl buffer (pH 8.0, 0.3% Triton-X). Reactions were conducted using 1 mL of spore suspension (OD_600_ = 1) with 0.8 mM substrates (C8, C12, and C16) in 1.7 mL microcentrifuge tubes. The reaction proceeded for 10 min at 42 °C with agitation (800 rpm) for each cycle. After reactions, the spores were collected by centrifugation (5000 g, 10 min), and the supernatant was decanted for colorimetric analysis. Tris-HCl buffer and fresh substrates were immediately added to the spore pellets to initiate the next cycle. Reaction conditions for all cycles were kept constant. Spores of TIED-LipB were recycled with the same procedure. To reuse TIED-APEX2 (OD_600_ = 1), each reaction cycle was performed with 125 *μ*M Amplex Red in PBS (pH 7.2) for 5 min. For reactions in methanol, spores of TIED-LipA in 1 mL PBS (OD_600_ = 1) were used. Before being subjected to reactions, the harvested spores in 1.7 mL microcentrifuge tubes were placed in a vacuum desiccator for 3 h to remove residual water within pellets. The dried spores were suspended in 1 mL of methanol (Fisher chemical, ≥ 99.9%) supplemented with 0.8 mM of C12 substrate. Each reaction was performed at 42 °C for 20 min before collecting spores by centrifugation (5000 g, 10 min). To renew spores with TIED-LipA, 2 *μ*L of spore suspension after the last reaction cycle was diluted in 5 mL of LB supplemented with the appropriate antibiotics. The starter culture was grown to saturation, and sporulation was induced as described in previous sections.

### Statistical analysis

All experimental data were plotted as mean ± s. e. m. unless otherwise mentioned. All enzymatic assays were performed with at least three biological replicates. The mean values determined from three biological replicates were fitted to the Michaelis Menten model for kinetics analyses. Error bars for *K*_*m*_ represent the 90% confidence interval (CI) of the fitting (Fig. 2i, 3g, and 4h). The thermal stability of lipases (Fig. 2l and 3j) was determined by fitting a Boltzmann sigmoid function to the experimental data. Statistical significance and *P* values were derived from two-tailed t-tests.

## ACKNOWLEDGEMENTS

This work is supported by start-up funds (S. S.) from the University of California, Irvine. The authors thank Professor David Tirrell for his helpful comments.

## AUTHOR CONTRIBUTIONS

Y. H. and S. S. designed experiments, analyzed the data, and wrote the manuscript. Y.H. and Z. C. performed the experiments and analyzed the data.

