## Supplementary Information for "Stress-tolerant, recyclable, and autonomously renewable biocatalyst platform enabled by engineered bacterial spores"

#### Table of Contents

1. **Genetic Construction** ........ S2
2. **Extended Data 1** ........ S4
3. **Extended Data 2** ........ S5
4. **Extended Data 3** ........ S6
5. **Extended Data 4** ........ S7
6. **Extended Data 5** ........ S8
7. **Extended Data 6** ........ S9
8. **Extended Data 7** ........ S10
9. **Extended Data 8** ........ S11
10. **Extended Data 9** ........ S12
11. **References** ........ S13

#### 1. Genetic construction

| Plasmid name | Description and Purpose | Transcriptional regulators | Output gene products | Tags (including linkers) | Integration site | Source |
|---|---|---|---|---|---|---|
| PBS1C-CotY-LipB-3XFLAG | TIED-LipB fused to CotY | $P_{T7}$, LacO | CotY-LipB-3XFLAG | 3XFLAG at C terminus; helical linker between CotY and LipB | amyE | This study |
| PBS1C-CotZ-LipB-3XFLAG | TIED-LipB fused to CotZ | $P_{T7}$, LacO | CotZ-LipB-3XFLAG | 3XFLAG at C terminus; helical linker between CotZ and LipB | amyE | This study |
| PBS1C-CotY-LipA-3XFLAG | TIED-LipA fused to CotY | $P_{T7}$, LacO | CotY-LipA-3XFLAG | 3XFLAG at C terminus; helical linker between CotY and LipA | amyE | This study |
| PBS1C-CotZ-LipA-3XFLAG | TIED-LipA fused to CotZ | $P_{T7}$, LacO | CotZ-LipA-3XFLAG | 3XFLAG at C terminus; helical linker between CotZ and LipA | amyE | This study |
| PBS1C-CotY-APEX2-3XFLAG | TIED-APEX2 fused to CotY | $P_{T7}$, LacO | CotY-APEX2-3XFLAG | 3XFLAG at C terminus; helical linker between CotY and APEX2 | amyE | This study |
| PBS1C-CotZ-APEX2-3XFLAG | TIED-APEX2 fused to CotZ | $P_{T7}$, LacO | CotZ-APEX2-3XFLAG | 3XFLAG at C terminus; helical linker between CotZ and APEX2 | amyE | This study |
| PBS4S-$P_{cotG}$-T7 RNAP | T7 RNAP expression | $P_{cotG}$ | T7 RNAP | | thrC | This study |
| PBS4S-$P_{cotV}$-T7 RNAP | T7 RNAP expression | $P_{cotV}$ | T7 RNAP | | thrC | This study |
| PBS4S-$P_{cotZ}$-T7 RNAP | T7 RNAP expression | $P_{cotZ}$ | T7 RNAP | | thrC | This study |

| | | | | | | |
|---|---|---|---|---|---|---|
| PET22b-6XHis-LipB-3XFLAG | LipB expression | $P_{T7}$, LacO | 6XHis-LipB-3XFLAG | 3XFLAG at C terminus 6XHis at N terminus | | This study |
| PET22b-6XHis-LipA-3XFLAG | LipA expression | $P_{T7}$, LacO | 6XHis-LipA-3XFLAG | 3XFLAG at C terminus 6XHis at N terminus | | This study |
| PET22b-6XHis-APEX2-3XFLAG | APEX2 expression | $P_{T7}$, LacO | 6XHis-APEX2-3XFLAG | 3XFLAG at C terminus 6XHis at N terminus | | This study |
| LacI-T7-sfgfp | T7 RNAP expression in vegetative *B. subtilis* cells | $P_{T7}$, LacO | sfGFP | | amyE | Tabor Lab[1] |
| PBS1C | Integration vector to amyE | | | | | BGSC ECE 257[2] |
| PBS4S | Integration vector to thrC | | | | | BGSC ECE 259[2] |

LacI-T7-sfgfp was obtained from Tabor lab[1]
PBS1C[2] and PBS4S[2] were obtained from the Bacillus Genetic Stock Center.

#### 2. Extended Data 1

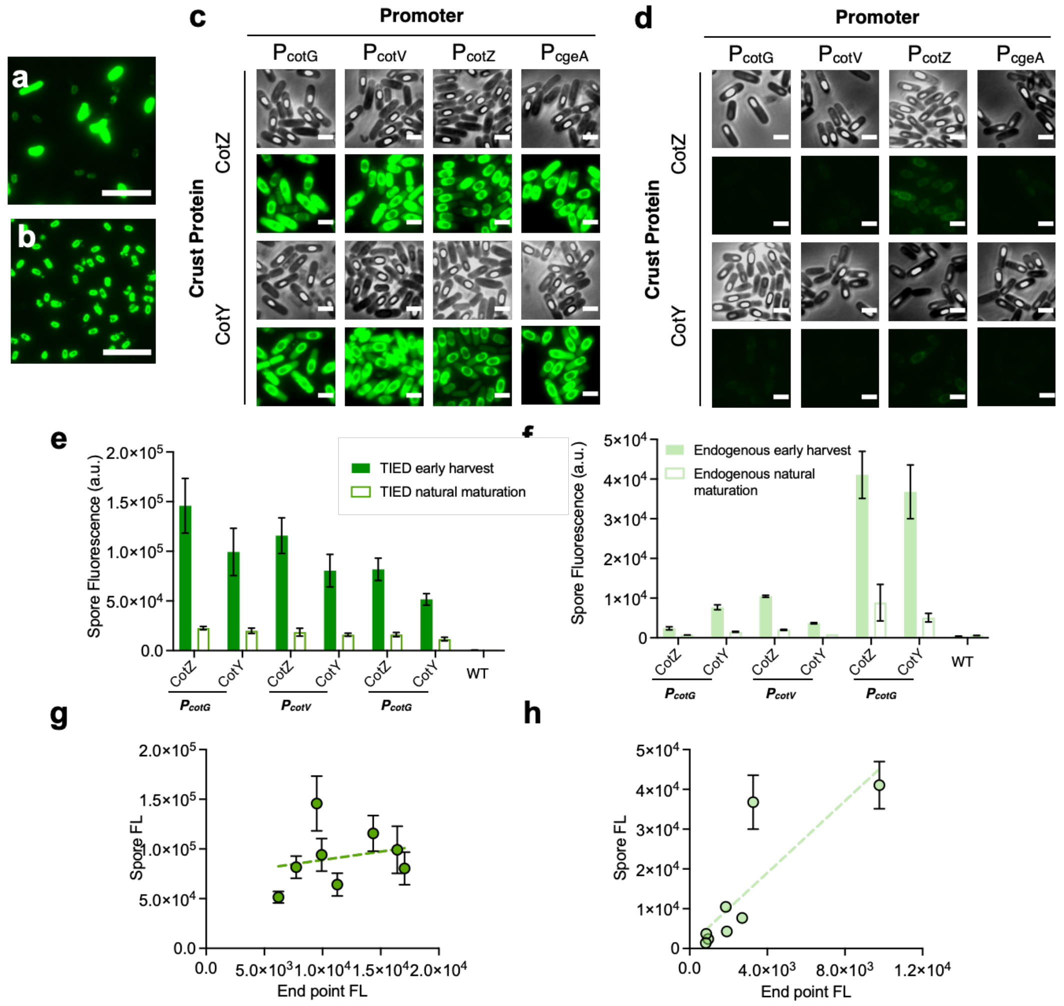


**Extended Data Figure 1.** **a**, A representative fluorescent microscopic image of TIED-mWasabi (combination of $P_{cotV}$ promoter and CotZ fusion partner) spores prepared by natural maturation after 24 h in SM medium. **b**, A representative fluorescent microscopic image of TIED-mWasabi (combination of $P_{cotV}$ promoter and CotZ fusion partner) spores prepared by early-harvest method with lysozyme digestion after growing in SM medium for 12 h. Same laser (3%) and exposure time (100 ms) were used for imaging. Scale bars, 10 $\mu$m. **c**, Representative fluorescent and phase-contrast microscopic images of sporulating cells of TIED-mWasabi variants before spores were harvested. **d**, Representative fluorescent and phase-contrast microscopic images of sporulating cells of endogenous constructs before spores were harvested. Scale bars, 2 $\mu$m. **e**, Purified TIED-mWasabi spores by the early harvest method (digested with 50 $\mu$g mL$^{-1}$ lysozyme) show higher retained spore fluorescence, relative to the natural maturation (n = 3 biological replicates). **f**, Purified spores of endogenous constructs by the early harvest method (digested with 50 $\mu$g mL$^{-1}$ lysozyme) show higher retained spore fluorescence, relative to the natural maturation. (n = 3 biological replicates). **g**, Bulk spore fluorescence of the TIED variants prepared through the early-harvest method as a function of total fluorescent protein expression. (n = 3 biological replicates. See Fig. 1b and 1c for endpoint fluorescence) Data were fitted with linear regression (dotted line). **h**, Bulk spore fluorescence of the endogenous counterparts prepared through the early-harvest method as a function of total fluorescent protein expression. Data were fitted with linear regression (dotted line). (n = 3 biological replicates).

#### 3. Extended Data 2

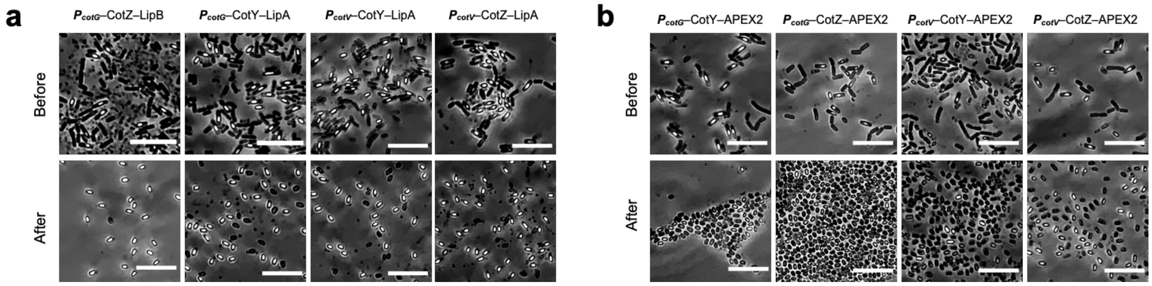


**Extended Data Figure 2.** **a**, Representative phase-contrast microscopic images of spores and cells of TIED-LipB and TIED-LipA variants. **b**, Representative phase-contrast microscopic images of spores and cells of TIED-APEX2 variants. Top panel: before lysozyme digestion. Bottom panel: after lysozyme digestion.

#### 4. Extended Data 3

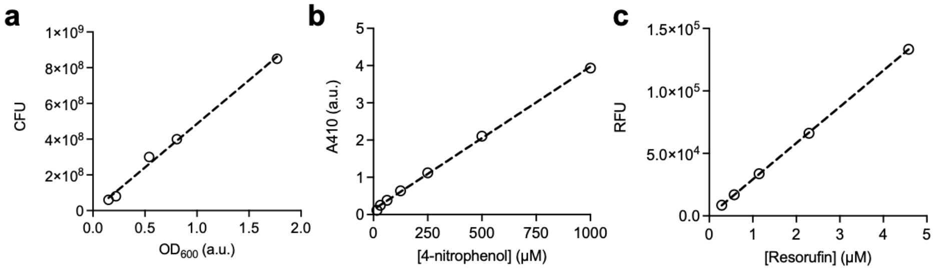


**Extended Data Figure 3.** **a**, $OD_{600}$–CFU standard calibration curve. **b,** Absorbance at 410 nm as a function of *p*-nitrophenol concentration. **c**, Relative fluorescence emission unit ($\lambda_{ex}/\lambda_{em}$ = 530/590 nm) as a function of sodium resorufin concentration.

#### 5. Extended Data 4

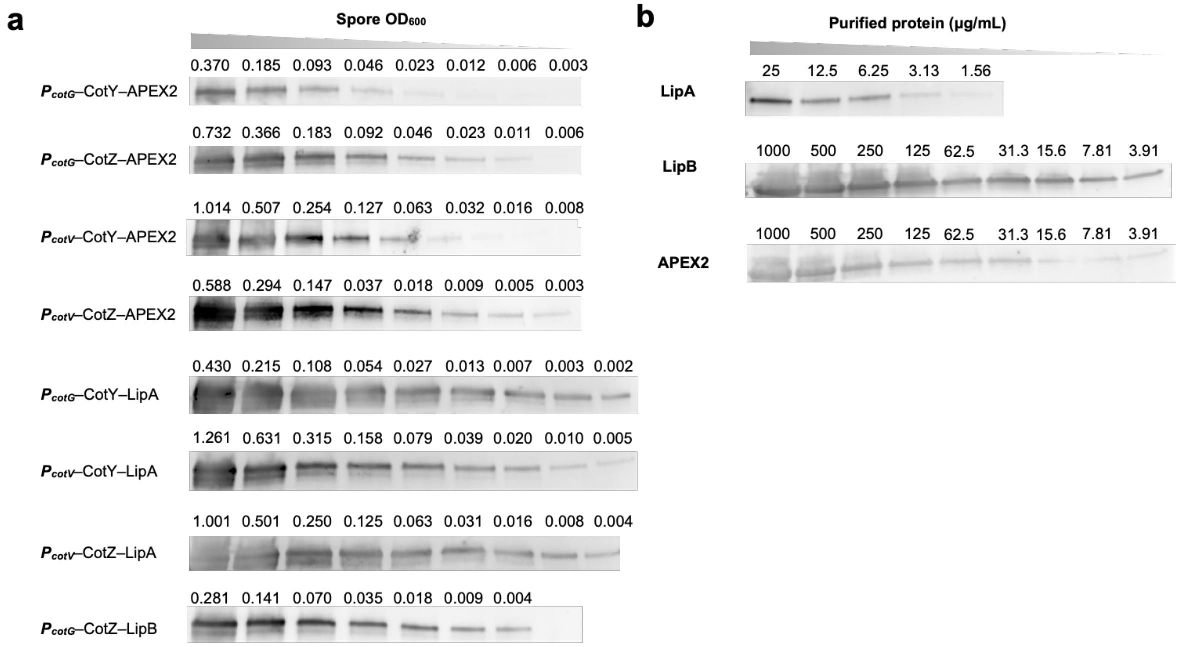


**Extended Data Figure 4.** **a**, Western blot images of lysates prepared from spores of TIED variants with the indicated $OD_{600}$. Bands correspond to the monomers of the respective fusion protein. **b**, Western blot images of solutions of purified enzymes (PBS) with the indicated concentrations. Bands correspond to the size of the respective enzyme.

#### 6. Extended Data 5

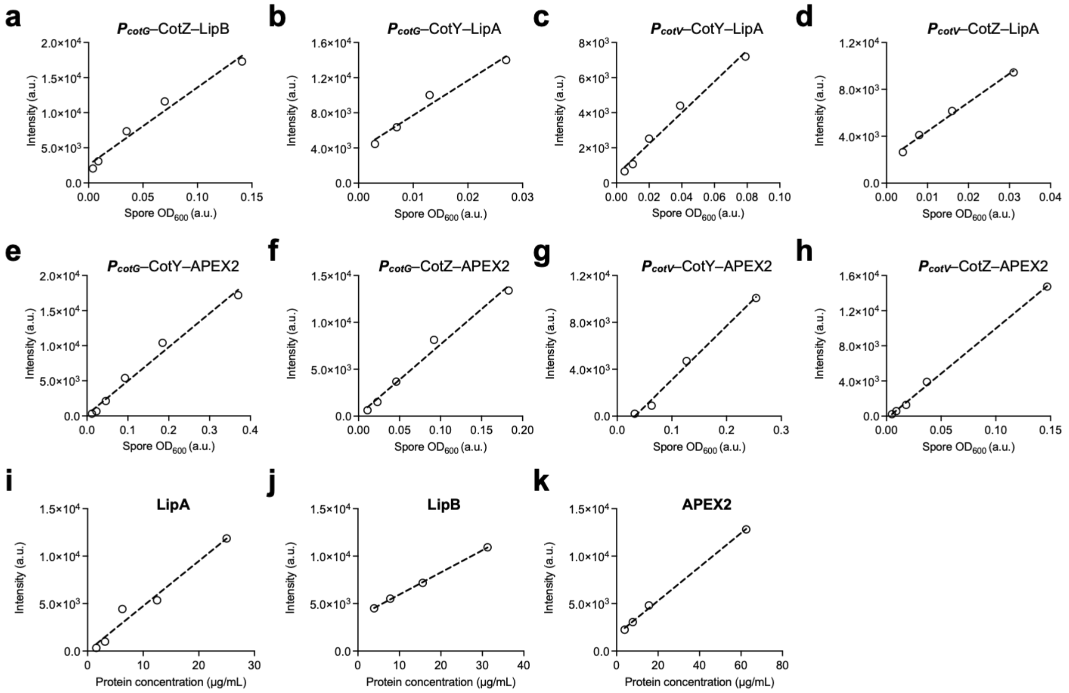


**Extended Data Figure 5. a-h**, Quantified band intensities of spore lysates as a function of spore $OD_{600}$. **i–k**, Quantified band intensities of purified free enzymes as a function of their concentration. Bands corresponding to the fusion proteins (Extended Data Fig. 4) were quantified with Image J and fitted with linear regression (dotted lines).

### 7. Extended Data 6

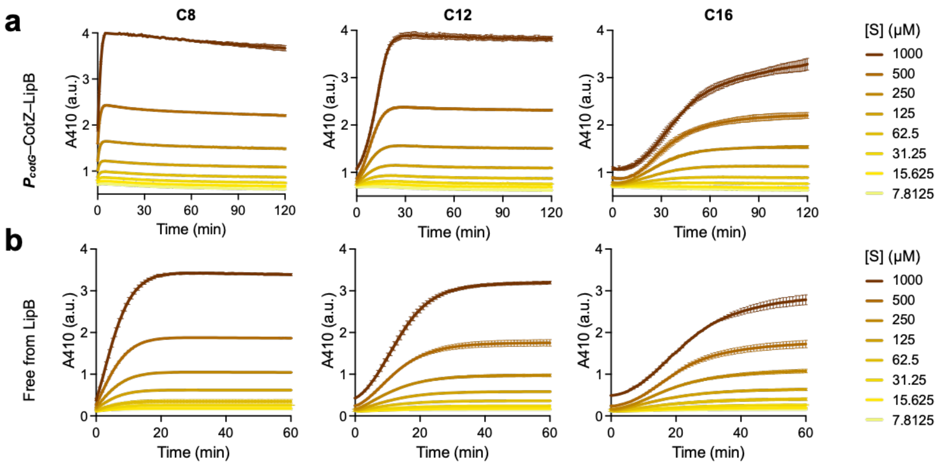


**c**

| Entry | Loading (enzyme/spore) | S | $V_{max}$ (μM/min) | $K_m$ (μM) |
|---|---|---|---|---|
| | | C8 | 379±47.1 | 1370±259 |
| $P_{cotG}$ CotZ–LipB | $1.75 \times 10^7$ | C12 | 80.6±5.76 | 1120±129 |
| | | C16 | 25.2±41.9 | 1090±294 |
| | | C8 | 152±17.0 | 1390±234 |
| Free form LipB | N/A | C12 | 91.5±19.4 | 1900±559 |
| | | C16 | 30.9±3.00 | 599±114 |

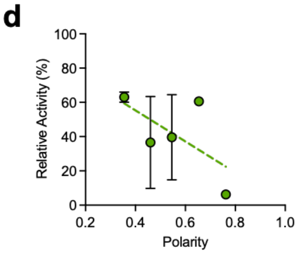


**Extended Data Figure 6.** **a**, TIED-LipB (combination of $P_{cotG}$ promoter and CotZ fusion partner, spore $OD_{600}$ of 0.5) catalyzed conversion of C8, C12, and C16 over time. The absorbance of the reaction product, *p*-nitrophenol, at 410 nm was monitored on a microplate reader (*n* = 3 biological replicates). **b**, Free form LipB (1 μg $mL^{-1}$) catalyzed conversion of C8, C12, and C16 over time (*n* = 3 biological replicates). **c**, The maximum rate of reaction ($V_{max}$) and the Michaelis Menten constant ($K_m$) of the reactions catalyzed by TIED-LipB and free form LipB. The constants were determined by fitting the rates of reaction at various substrate concentrations with the Michaelis Menten model. **d**, Relative activities of free form LipB in polar organic solvents (methanol, ethanol, 2-propanol, acetone, and acetonitrile) as a function of solvent polarity. The dotted line indicates fitting with linear regression (*n* = 3 biological replicates).

#### 8. Extended Data 7

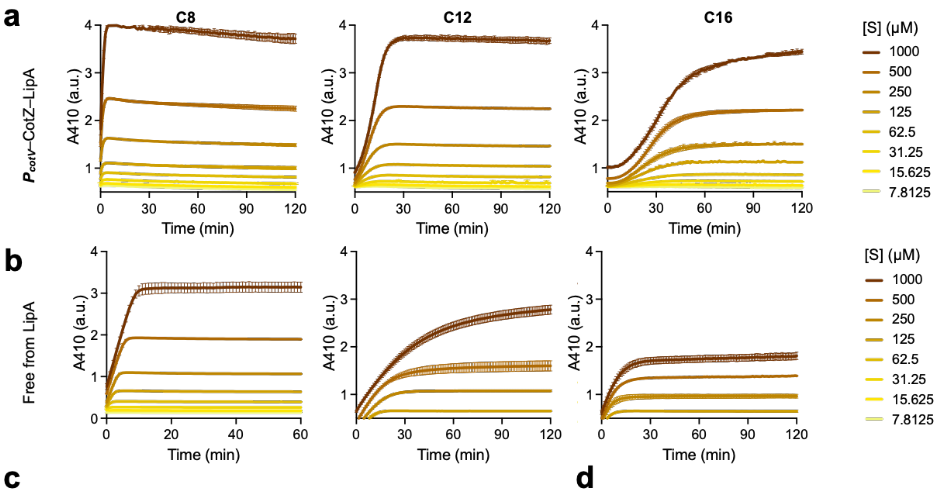


c

| Entry | Loading (enzyme/spore) | S | $V_{max}$ (μM/min) | $K_m$ (μM) |
|---|---|---|---|---|
| $P_{cotV}$ CotZ–LipA | 2.93 × 10[7] | C8 | 511±90.4 | 1490±392 |
| | | C12 | 129±0.860 | 1890±178 |
| | | C16 | 30.0±3.57 | 926±188 |
| Free form LipA | N/A | C8 | 84.1±17.0 | 163±35.5 |
| | | C12 | 16.2±1.34 | 86.1±25.4 |
| | | C16 | 26.4±3.02 | 69.6±19.1 |

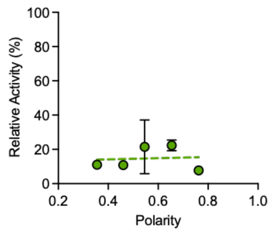


**Extended Data Figure 7. a**, TIED-LipA (combination of $P_{cotV}$ promoter and CotZ fusion partner, spore $OD_{600}$ of 0.5) catalyzed conversion of C8, C12, and C16 over time. The absorbance of the reaction product, *p*-nitrophenol, at 410 nm was monitored on a microplate reader (*n* = 3 biological replicates). **b**, Free form LipA (1 μg $mL^{-1}$) catalyzed conversion of C8, C12, and C16 over time (*n* = 3 biological replicates). **c**, The maximum rate of reaction ($V_{max}$) and the Michaelis Menten constant ($K_m$) of the reactions catalyzed by TIED-LipA and free form LipA. The constants were determined by fitting the rates of reaction at various substrate concentrations with the Michaelis Menten model. **d**, Relative activities of free form LipA in polar organic solvents (methanol, ethanol, 2-propanol, acetone, and acetonitrile) as a function of solvent polarity. The dotted line indicates fitting with linear regression (*n* = 3 biological replicates).

#### 9. Extended Data 8

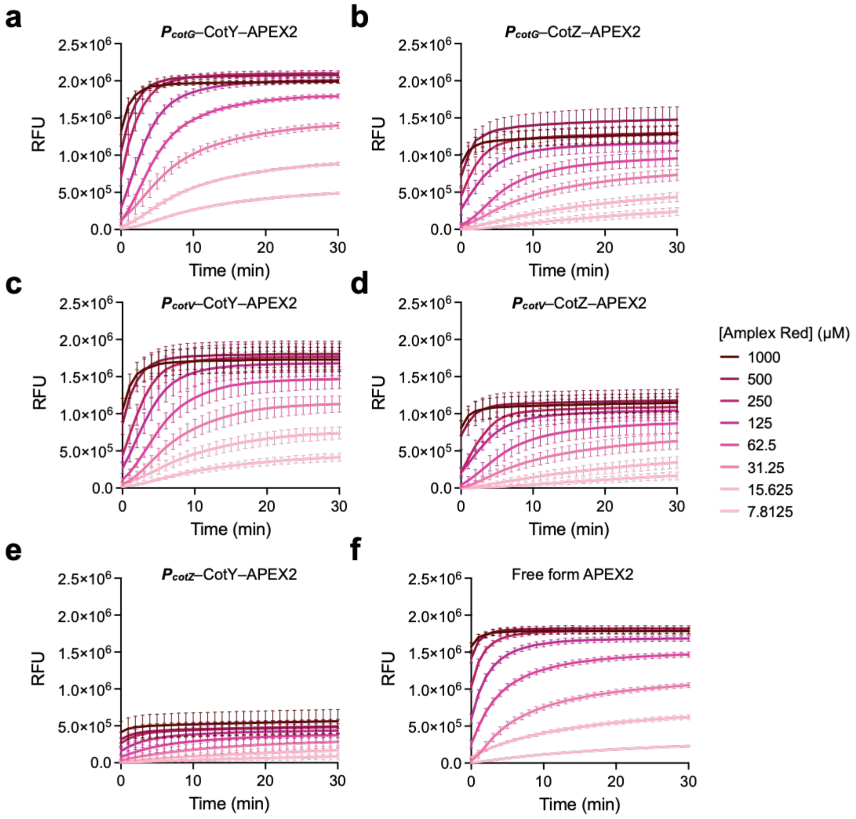


**g**

| Entry | Loading (enzyme/spore) | $V_{max}$ (μM/min) | $K_m$ (μM) |
|---|---|---|---|
| $P_{cotG}$ CotY–APEX2 | $6.75 \times 10^6$ | 17.5±3.33 | 108±45.2 |
| $P_{cotG}$ CotZ–APEX2 | $1.23 \times 10^7$ | 12.4±3.65 | 174±95.6 |
| $P_{cotV}$ CotY–APEX2 | $2.85 \times 10^6$ | 13.67±3.36 | 132±66.4 |
| $P_{cotV}$ CotZ–APEX2 | $1.71 \times 10^7$ | 9.28±3.61 | 143±111 |
| Free form APEX2 | N/A | 14.8±2.51 | 93±36.3 |

**Extended Data Figure 8.** **a**, Conversion of Amplex Red to resorufin over time by TIED-APEX2 with the combination of $P_{cotG}$ promoter and CotY fusion partner. **b**, Conversion of Amplex Red to resorufin over time by TIED-APEX2 with the combination of $P_{cotG}$ promoter and CotZ fusion partner. **c**, Conversion of Amplex Red to resorufin over time by TIED-APEX2 with the combination of $P_{cotV}$ promoter and CotY fusion partner. **d**, Conversion of Amplex Red to resorufin over time by TIED-APEX2 with the combination of $P_{cotV}$ promoter and CotZ fusion partner. **e**, Conversion of Amplex Red to resorufin over time by TIED-APEX2 with the combination of $P_{cotZ}$ promoter and CotY fusion partner. **f**, Conversion of Amplex Red to resorufin over time by free form APEX2 (1 μg $mL^{-1}$). TIED-APEX2 with spore $OD_{600}$ of 0.1 was used for each reaction. **g**, The maximum rate of reaction ($V_{max}$) and the Michaelis Menten constant ($K_m$) of the reactions catalyzed by TIED-APEX2 and free form APEX2. The constants were determined by fitting the rates of reaction at various substrate concentrations with the Michaelis Menten model.

#### 10. Extended Data 9

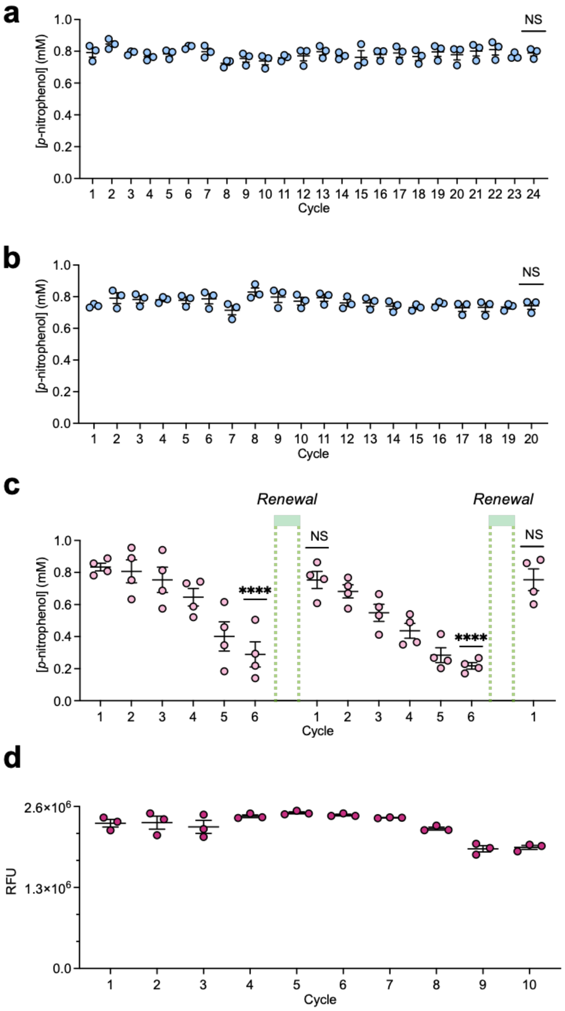


**Extended Data Figure 9. a**, Recycling TIED-LipA (combination of $P_{cotV}$ promoter and CotZ fusion partner, spore $OD_{600}$ of 1.0) for conversion of C8 (0.8 mM) in Tris-HCl (50 mM, pH 8.0). **b**, Recycling TIED-LipA (combination of $P_{cotV}$ promoter and CotZ fusion partner, spore $OD_{600}$ of 1.0) for conversion of C12 (0.8 mM) in Tris-HCl. **c**, Recycling and renewal of TIED-LipB (combination of $P_{cotG}$ promoter and CotZ fusion partner, spore $OD_{600}$ of 1.0) for conversion of C16 (0.8 mM) in Tris-HCl. **d**, Recycling TIED-APEX2 (combination of $P_{cotG}$ promoter and CotY fusion partner, spore $OD_{600}$ of 1.0) for conversion of Amplex Red (0.125 mM) in PBS (pH 7.2). *P* values were determined by two-tail t-tests against the pristine activity of TIED-enzymes ($1^{st}$ cycle). $^{*}P < 0.05$, $^{**}P < 10^{-2}$, $^{***}P < 10^{-3}$, $^{****}P < 10^{-4}$.
